# High-Recovery Desalting Tip Columns for a Wide Variety of Peptides in Mass Spectrometry-based Proteomics

**DOI:** 10.1101/2024.07.19.604362

**Authors:** Eisuke Kanao, Shunsuke Tanaka, Ayana Tomioka, Tetsuya Tanigawa, Takuya Kubo, Yasushi Ishihama

## Abstract

In mass spectrometry-based proteomics, loss-minimized peptide purification techniques play a key role in improving sensitivity and coverage. We have developed a desalting tip column packed with thermoplastic polymer-coated chromatographic particles, named ChocoTip, to achieve high recoveries in peptide purification by pipette-tip-based LC with centrifugation (tipLC). ChocoTip identified more than twice as many peptides from 20 ng of tryptic peptides from Hela cell lysate compared to a typical StageTip packed with chromatographic particles entangled in a Teflon mesh in tipLC. The high recovery of ChocoTip in tipLC was maintained for peptides with a wide variety of physical properties over the entire retention time range of the LC/MS/MS analysis, and was especially noteworthy for peptides with long retention times. These excellent properties are attributable to the unique morphology of ChocoTip, in which the thermoplastic polymer covers the pores, thereby inhibiting irreversible adsorption of peptides into mesopores of the chromatographic particles.

ChocoTip is expected to find applications, especially in clinical proteomics and single-cell proteomics, where sample amounts are limited.

## INTRODUCTION

In the realm of biological research, mass spectrometry (MS)-based proteomics is indispensable for the identification, quantification, and characterization of proteins that play key roles in maintaining physiological cellular functions.^1–4^ Bottom-up proteomics is the workhorse for MS-based proteomics, constituting a well-defined multistep process that integrates various methodologies and instrumentation.^5,6^ The process can be broadly categorized into three main steps: sample preparation, liquid chromatography/tandem MS (LC/MS/MS) analysis, and database search.^7^ The characteristics and limitations of each step in the proteomic process significantly influence the data quality. Because errors and biases introduced during the initial sample preparation step can propagate throughout the entire experiment,^8,9^ the success of bottom-up proteomics depends upon optimal and consistent sample preparation.

Purifying and concentrating digested proteins is a key part of the sample preparation process, with solid-phase extraction (SPE) being the mainstream technique.^10–12^ SPE involves either centrifugation or vacuum suction through hydrophobic reversed-phase (RP) materials, employing alternating mobile phases for the trapping, washing, and elution steps. This process, often referred to as ’desalting,’ primarily aims to remove buffer salts.^13^ These salts can interfere with the ionization process and reduce the lifespan of the analytical system. The Stop-and-Go Extraction Tip (StageTip) is one of the most common microcolumns utilized as an offline desalting system, comprising a small disk of hydrophobic particles entangled in a Teflon mesh within a pipette tip.^14,15^ In contrast to online systems, StageTip offers the advantage of circumventing limitations imposed by the size and characteristics of trap columns and downstream analytical columns. Furthermore, StageTip can process multiple peptide samples simultaneously, thereby minimizing sample preparation time. StageTip can also prevent the carryover of peptides into LC/MS/MS due to the disposable format.

Recent advancements in this field include in-StageTip^16,17^ and the On-microSPE method^18,19^ utilizing StageTips as solid-phase reactors throughout the sample preparation process, covering steps from cell lysis to peptide purification. These approaches effectively minimize contamination and sample loss in the overall workflow. Another advancement is the Evosep One system, which seamlessly integrates StageTip directly into the downstream LC/MS/MS workflow.^20,21^ This integration is designed to increase throughput and robustness, especially in applications related to single-cell proteomics and clinical proteomics. Nevertheless, the desalting step still poses a significant challenge, with the risk of losing peptide samples due to inadequate retention or irreversible adsorption onto hydrophobic RP materials.^22,23^ Traditionally, hydrophobic ion-pair reagents^24,25^ or chemical modifications of peptides^26–28^ have been employed to improve the peptide retention on RP materials. However, these chemical approaches may have the drawback of reducing MS sensitivity, owing to a decline in electrospray ionization efficiency^29–31^ or peptide losses within the complex workflow.^32,33^ Porous graphite carbon (PGC) has been used as a hydrophobic RP material to enhance the recovery of hydrophilic peptides through strong hydrophobic and π interactions.^34–37^ However, the overly strong intermolecular interactions with PGC resulted in loss of hydrophobic peptides.^37,38^ We previously introduced CoolTip, where the recovery efficiency of hydrophilic peptides was increased by cooling the StageTip during the desalting step.^39^ Still, there appeared to be significant potential for improvement in the material design of StageTip to further enhance sensitivity and coverage in MS-based proteomics.

Here, we present a unique hybrid polymer designed to improve the performance of StageTip. The polymer was synthesized by thermally kneading commercially available hydrophobic particles of styrene-divinylbenzene copolymer (St-DVB) into ethylene vinyl acetate copolymer (EVA) thermoplastic resin. The EVA forms a flexible sponge-like monolithic carrier with μm-sized through-pores (SPongy Monolith; SPM), functioning simultaneously as the column material and frit.^40–42^ The through-pores facilitate low-pressure chromatographic separation, ensuring the efficient processing of biological samples without clogging. The St- DVB particles embedded in the surface of the monolithic carrier enhance peptide retention through strong hydrophobic interaction. A pipette tip containing this hybrid polymer, named ChocoTip (CHrOmatographic particles COated by a Thermoplastic polymer Immobilized in Pipette tip), was developed to address the sample loss issues in conventional StageTip methods. The simplicity and outstanding peptide recovery efficiency of ChocoTip suggest its potential suitability as a universal platform for peptide purification in ultra-sensitive proteomics applications.

## MATERIALS AND METHODS

### Materials

UltraPure Tris Buffer was purchased from Thermo Fisher Scientific (Waltham, MA). Sequencing-grade modified trypsin was purchased from Promega (Madison, WI). Water was purified by a Millipore Milli-Q system (Bedford, MA). Empore SDB-XC disks and InertSep PLS-2 were purchased from GL Sciences (Tokyo, Japan). Polyethylene frit was purchased from Agilent Technologies (Santa Clara, CA). Protease inhibitors were purchased from Sigma-Aldrich (St. Louis, MO). Blunt-end 16- or 17-gauge syringe needles were purchased from Hamilton (Reno, NV). 200 µL pipette tips were purchased from Gilson (Middleton, WI) and used for the preparation of StageTips. All other chemicals and reagents were purchased from Fujifilm Wako (Osaka, Japan) unless otherwise specified.

### Preparation of SPM-tip and ChocoTip

SPM-tip and ChocoTip were prepared in a similar manner to the previously reported method.^40,41^ Briefly, 37□wt% of polyolefin chips containing 15 % vinyl acetate, 55□wt% of pore templates (pentaerythritol), and 8 wt% of auxiliary pore templates (poly(oxyethylene/ oxypropylene) triol) were blended at 130□°C and homogeneously kneaded.^40,41^ For ChocoTip, St-DVB particles (mean particle diameter; 70□μm) were added at 20 wt% to SPM during the kneading step.^42,43^ The resulting material was extruded at 130 °C, and the resulting string-shaped material was immediately cooled in water for solidification. The product was washed with water using ultrasonication to remove water-soluble compounds. The porosity of the obtained hybrid material was about 75 % and the diameter of the cross-section across its entire length was 1.5 mm. The string-shaped material was then sliced at intervals of 2.0 mm (SPM pellets). Due to the elasticity of the SPM, the packing procedure is simple (Figure 1a). The pellets were immersed in water and thoroughly wetted, and pushed straight into the 200 µL pipette tip using a 17-gauge syringe needle. The pellets were carefully inserted to prevent distortion or wrinkling inside the cartridge. Then the prepared tip was washed with methanol (200 μL×5) and water (200 μL×1) to remove the pore templates and homogenize the packing state. Morphology observation of SPM-tip and ChocoTip was carried out using a field-emission scanning electron microscope (SEM; JSM6700-M, JEOL, Tokyo, Japan).

**Figure 1.**
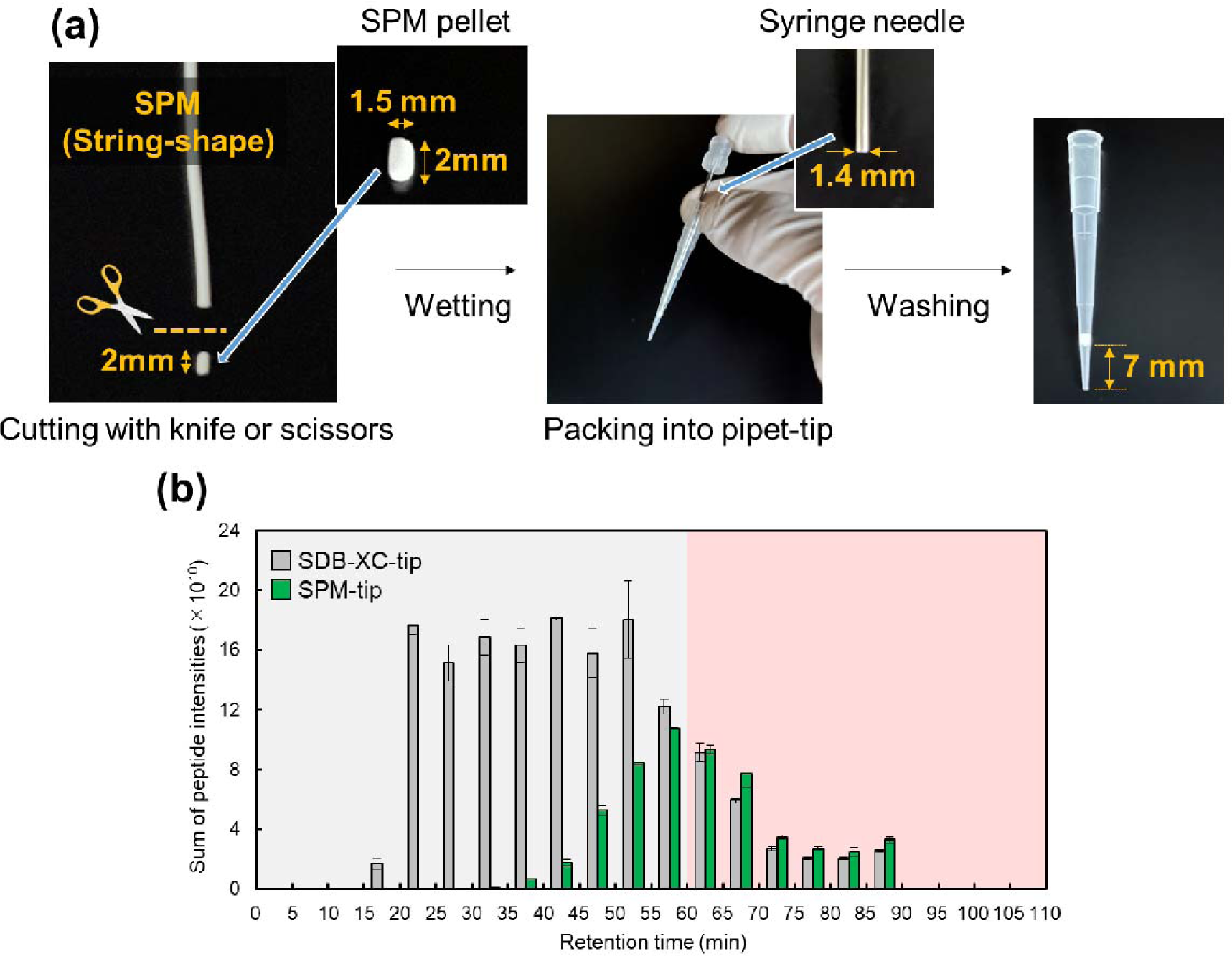
Preparation process and desalting performance of SPM-tip. (a) Preparation process of SPM-tip. (b) Comparison of the sum of peptide intensities binned by retention time between SPM-tip and SDB-XC-tip. 500 ng of tryptic peptides from HeLa cell lysates were loaded onto each StageTip, and a sample equivalent to 250 ng of peptides was injected into the LC/MS/MS system. The error bars indicate the SDs of triplicate analyses with three StageTips.

### Cell Culture

HeLa S3 cells obtained from the JCRB Cell Bank (Osaka, Japan) were cultured to 80% confluency in Dulbecco’s modified Eagle’s medium containing 10% fetal bovine serum in 10- cm diameter dishes. Cells were washed twice with ice-cold PBS, collected using a cell scraper, and pelleted by centrifugation.

### Protein Extraction

HeLa cell lysates were digested using phase-transfer surfactant–aided trypsin digestion as described previously.^43,44^ Briefly, the cell pellets were suspended in 1 mL of buffer (12 mM sodium deoxycholate, 12 mM sodium lauroyl sarcosinate in 100 mM Tris-HCl, pH 9.0) containing protease inhibitors. The cells were incubated on a heating block at 95 °C for 5 min and then sonicated for 20 min. The extracted proteins were quantified with a BCA protein assay kit, reduced with 10 mM dithiothreitol for 30 min, and alkylated with 50 mM iodoacetamide for 30 min in the dark. The samples were diluted 5-fold with 50 mM ammonium bicarbonate and then digested with Lys-C for 3 h at room temperature and with trypsin overnight at 37 °C. 1 mL of ethyl acetate was added to 1 mL of the digested solution, and the mixture was acidified with 0.5% trifluoroacetic acid (TFA) (final concentration). The samples were vortexed for 2 min and centrifuged at 15,800g for 2 min to completely separate the aqueous and organic phases. The aqueous phase was collected, dried, resuspended in 0.1% TFA/5% acetonitrile (ACN) solution, and desalted as follows.

### Desalting by StageTip

SPM-tip and ChocoTip were prepared as described above. For comparison, SDB-XC-tips were manufactured by stamping out three pieces or a single piece of Empore SDB-XC disk with a 16-gauge syringe needle and packing into 200 µL pipette tips. Desalting procedures are summarized in Table 1, and were in line with those in the previous study.^39^ After desalting, peptides were dried in a vacuum centrifuge and the residue was dissolved in 4% ACN with 0.5% TFA. Throughout this study, the sample loading amount into StageTip was referenced to the protein amount before the digestion process, as determined by BCA assay.

**Table 1.** Summary of solvents used for peptide desalting.

| Step number | Content | Solution | Centrifugation conditions |
| --- | --- | --- | --- |
| 1 | Washing | 80% ACN and 0.1% TFA | 1000 g x 2 min |
| 2 | Equilibration | 4% ACN and 0.1% TFA | 1000 g x 2 min |
| 3 | Sample loading | 4% ACN and 0.1% TFA | 800 g x 3 min |
| 4 | Washing | 4% ACN and 0.1% TFA | 1000 g x 2 min |
| 5 | Elution | 80% ACN and 0.1% TFA | 800 g x 3 min |

### LC/MS/MS analysis

Orbitrap system

Unless otherwise described, LC/MS/MS analysis was carried out in the data-dependent acquisition (DDA) mode with a FAIMS Pro Duo interface connected to an Orbitrap Fusion Lumos Tribrid MS. The electrospray voltage was set to 2.4 kV in the positive mode. The FAIMS mode was set to standard resolution, and the total carrier gas flow was 4.6 L/min. The compensation voltage (CV) was set to -40, -60, and -80, and the cycle time of each CV experiment was set to 1 s. The mass range of the survey scan was from 300 to 1,500 *m/z* with a resolution of 120,000, standard automatic gain control (AGC), and a maximum injection time of 50 ms. The MS/MS scan was performed using an ion trap with a rapid ion trap scan rate, standard AGC, a maximum injection time of 35 ms, and an isolation window of 1.6 *m/z*. The precursor ions were fragmented by higher-energy collisional dissociation with a normalized collision energy of 30%. The exclusion duration was set to 20 s. LC was performed on an Ultimate 3000 pump (Thermo Fisher Scientific) and an HTC-PAL autosampler (CTC Analytics) using self-pulled needle columns (150 mm length, 100 μm ID, 6 μm needle opening) packed with Reprosil-Pur 120 C18-AQ 1.9 μm reversed-phase material (Dr. Maisch, Ammerbuch, Germany).^45^ The injection volume was 5 μL, and the flow rate was 500 nL/min. Separation was achieved by applying a three-step linear gradient of 4−8% ACN in 5 min, 8−32% ACN in 60 min, 32−80% ACN in 5 min, and 80% ACN for 10 min in 0.5% acetic acid.

### Q-TOF system

For comparison, we analyzed the same samples by LC/TIMS/Q/TOF using a timsTOF Pro 2 (Bruker, Bremen, Germany). The TIMS section was operated with a 100 ms ramp time and a scan range of 0.6-1.5 V s cm^-2^. One cycle was composed of 1 MS scan followed by 10 parallel accumulation serial fragmentation MS/MS scans. MS and MS/MS spectra were recorded from *m/z* 100 to 1,700. A polygon filter was applied to avoid selecting singly charged ions. The quadrupole isolation width was set to *m/z* 2 or 3. The collision energy was ramped stepwise as a function of increasing ion mobility: 42 eV for 0-6% of the ramp time; 32 eV from 6-22%; 37 eV from 22-44%; 42 eV from 44-67%; 47 eV from 67-89%; and 51 eV for the remainder. The timsTOF Pro 2 was connected to the same LC system and autosampler as the Orbitrap Fusion Lumos Tribrid MS using self-pulled needle columns (250 mm length, 100 μm ID) packed with Reprosil-Pur 120 C18-AQ 1.9 μm reversed-phase material (Dr. Maisch, Ammerbuch, Germany). The injection volume was 5 μL, and the flow rate was 500 nL/min. Separation was achieved by applying a three-step linear gradient of 4−8% ACN in 5 min, 8−32% ACN in 60 min, 32−80% ACN in 5 min and 80% ACN for 10 min in 0.1% formic acid.

### Database Searching and Data Processing

The raw MS data was analyzed by MaxQuant (MQ) version 1.6.17.0.^46^ Peptides and proteins were identified by an automated database search using Andromeda against the human SwissProt Database (version 2022-10, 20,401 protein entries). The data files collected from FAIMS experiments were split into a set of MaxQuant-compliant MzXML files using FAIMS MzXML Generator (https://github.com/coongroup/FAIMS-MzXML-Generator). For data files of the Orbitrap system, the precursor mass tolerance of 20 ppm for the first search, 4.5 ppm for the main search, and the fragment ion mass tolerance of 0.5 Da were set. For the Q-TOF system, the precursor mass tolerance of 20 ppm for the first search, 10 ppm for the main search, and the fragment ion mass tolerance of 40 ppm were set. The enzyme was set as Trypsin/P with two missed cleavages. “Cysteine carbamidomethylation” was set as a fixed modification and “methionine oxidation” and “acetylation on the protein N-terminus” were set as variable modifications. The search results were filtered with FDR <1% at the peptide spectrum match (PSM) and protein levels. The match-between-runs algorithm (MBR) was utilized through the “Identification” subtab in the “Global Parameters” tab of MaxQuant to mitigate the missing value problem. The default settings for MBR were used (0.7 min match window and 20 min alignment time). Proteins that have “Only identified by site”, “potential contaminants” and “reverse sequences” were removed for data analysis.

## RESULTS AND DISCUSSION

We first compared SPM-tip with SDB-XC-tip for desalting 500 ng of tryptic peptides from HeLa cell lysates. The preparation of SPM-tip is illustrated in Figure 1a. In brief, a string-shaped SPM with a diameter of 1.5 mm was cut into pellets of 2.0 mm in length using a scalpel or scissors. The pellets were inserted straight into a 200 μL pipette tip and gently pressed with a 17-gauge stainless steel needle (1.4 mm o.d.) until the needle became caught on the inner wall, preventing further insertion. The sponge-like flexibility allowed the SPM to fit snugly into the pipette tip. On the other hand, SDB-XC-tips were prepared by stamping Empore SDB-XC disks with a 16-gauge needle (1.6 mm inner diameter) according to the original method for preparing StageTips using Empore disks.^15^ Three Empore disks were stacked in order to keep the volume of the stationary phase roughly the same as that of the SPM-tip, according to the previous report.^39^

A SEM image of a cross-section of SPM-tip is shown in Figure S1. Desalting steps were rapidly completed with simple centrifugation due to the high permeability of the SPM-tip. After desalting, a sample equivalent to 250 ng of peptides was injected onto the LC/MS/MS. Figure 1b shows the sum of peptide intensities based on the peptide-ion intensity of MQ label-free quantification binned by retention time in LC/MS/MS, with each retention time window partitioned by 5 minutes for both SPM- and SDB-XC-tip. The sum of peptide intensities was higher for SDB-XC-tip, and SPM-tip exhibited a limited ability to cature hydrophilic peptides with short retention times (Figure 1b, gray region). The weak retention of peptides on SPM-tip could be attributed to the less hydrophobic structure compared to St- DVB particles in the Empore SDB-XC disk. Interestingly, the sum of intensities of peptides with long retention times was similar or slightly higher in the case of SPM-tip (Figure 1b, red region). This result strongly suggested that St-DVB particles and SPM are suitable for the retention of peptides of different hydrophobicity and that hybridization of these materials may allow comprehensive recovery of peptides.

Then, we prepared ChocoTip to compare the desalting performance with that of SDB-XC-tip. The packing process for ChocoTip employed the same procedure used for SPM-tip. Figure 2a shows the sum of peptide intensities for ChocoTip and SDB-XC-tip. As expected, both the sum of peptide intensities and the number of identified peptides were higher for ChocoTip compared to SDB-XC-tip, especially in the long retention time region (Figure 2a, Figure S2a). Notably, the excellent recovery efficiency of ChocoTip was highlighted with a smaller sample amount. After desalting 20 ng tryptic peptides from HeLa cell lysates using each StageTip, a sample equivalent to 10 ng of peptides was injected into the LC/MS/MS system. In ChocoTip, the sum of peptide intensities and the number of identified peptides were significantly higher than in the case of SDB-XC-tip over the entire retention time range (Figure 2b, Figure S2b). Specifically, while 1158 unique peptides were identified using SDB-XC-tip and 4062 peptides were commonly identified, 6835 unique peptides were identified using ChocoTip (Figure 2c, Figure S2). The unique peptides identified in ChocoTip tended to be longer than the commonly identified peptides and unique peptides in SDB-XC-tip (Figure S3), suggesting that ChocoTip might be useful for bottom-up proteomics using long peptides, such as in post-translational modification analysis^47^ and structural proteomics using limited proteolysis,^48^ as well as middle-down proteomics.^49^ Furthermore, the peptide intensity of each commonly identified peptide was consistently higher for ChocoTip over the entire retention time range (Figure 2d) and the trend toward greater intensity was more pronounced at longer retention times. LC/TIMS/Q/TOF analysis was also carried out for peptides desalted with ChocoTip and SDB-XC-tip, and the results suggest the advantage could be independent of the downstream LC/MS/MS system (Figure S4). In brief, the ChocoTip method was demonstrated to be a suitable sample treatment for ultra-sensitive proteomics, giving superior results to the standard StageTip.

**Figure 2.**
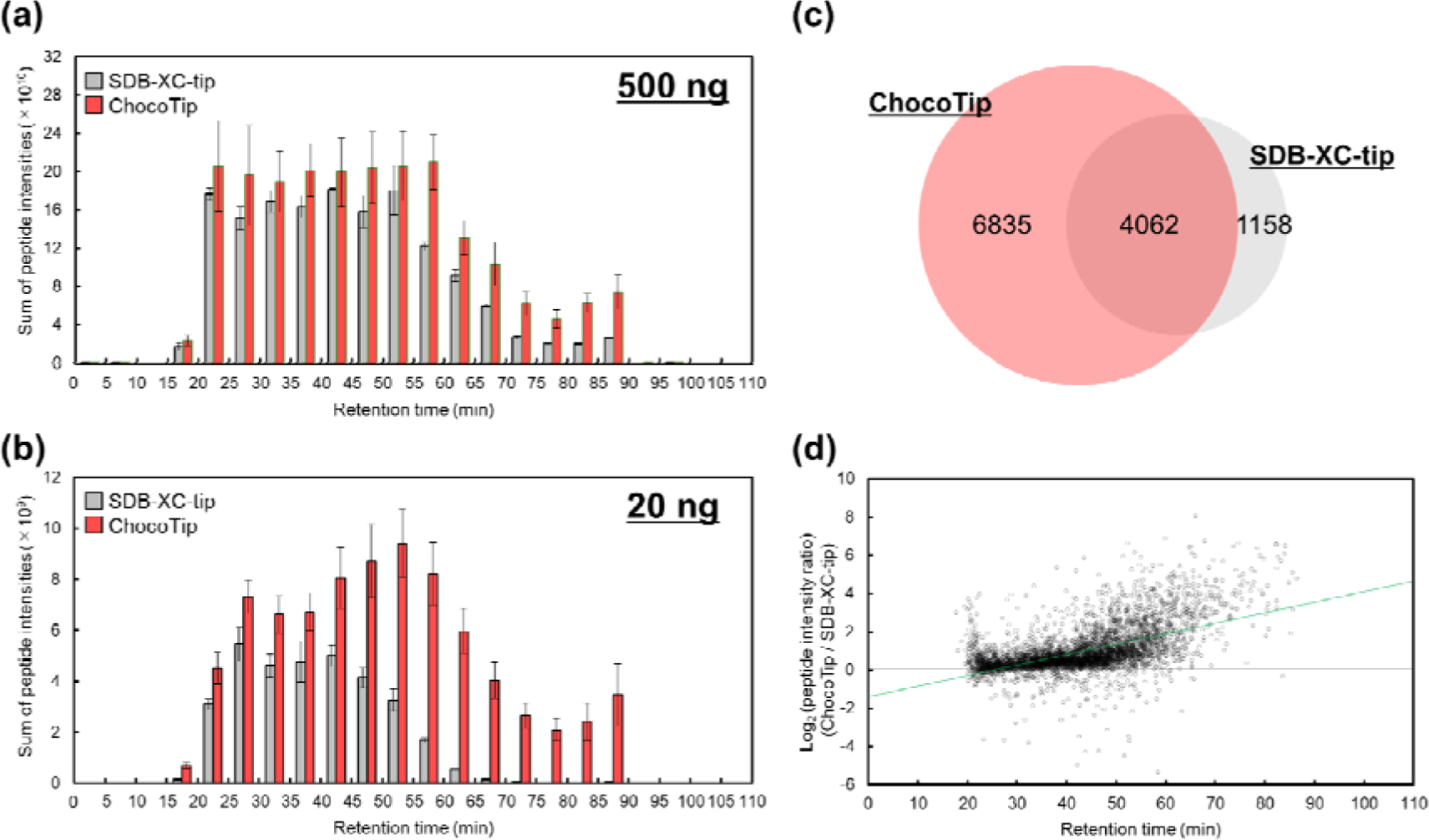
Desalting performance of ChocoTip compared to SDB-XC-tip. **(a),** (b) Comparison of the sum of peptide intensities binned by retention time between ChocoTip and SDB-XC-tip. StageTips were used for desalting (a) 500 ng or (b) 20 ng of tryptic peptides from HeLa cell lysates. A sample equivalent to (a) 250 ng or (b) 10 ng of peptides was injected into the LC/MS/MS system. The error bars indicate the SDs of triplicate analyses with three StageTips. (c) Overlap of identified peptides between ChocoTip and SDB-XC-tip. Only peptides identified in all triplicate analyses were used for the evaluation. (d) Scatter plot of the commonly identified peptide intensity ratio with ChocoTip and SDB-XC-tip versus the retention time. The green line is the linear regression line obtained by the least-squares method. Figure 2c and 2d were analyzed using the same LC/MS/MS datasets as Figure 2b.

To investigate the factors contributing to the highly efficient recovery of peptides with ChocoTip, we observed the surface morphology of ChocoTip with a scanning electron microscope (SEM) (Figure 3a). In ChocoTip, the surface of the St-DVB particles was partially enveloped by the fibrous structure of the monolithic carrier, and the particles were arranged on the surface of the μm-sized through-pores. This result suggested that the enhanced hydrophobicity of the surface of ChocoTip with St-DVB particles is the reason for the comprehensive retention of peptides. In addition, mercury intrusion porosimetry analysis was carried out to characterize the pore size distribution of the SDB-XC-tip, SPM-tip, and ChocoTip (Figure 3b). In each material, μm-sized through-pores were observed. Interestingly, while nm-sized mesopores of St-DVB particles were observed in the SDB-XC-tip, they were not observed in ChocoTip, even though it contains St-DVB particles. This could be attributed to thermally melted EVA filling the mesopores of St-DVB particles during the mixing process. The mesopores of RP hydrophobic materials often irreversible adsorb peptides, decreasing the recovery efficiency in the purification process.^50^ Thus, the high recovery in the case of ChocoTip could be a consequence of the enhanced hydrophobicity of the through-pore surface of SPM with embedded St-DVB particles, as well as the suppression of irreversible peptide adsorption at the mesopores of St-DVB particles.

**Figure 3.**
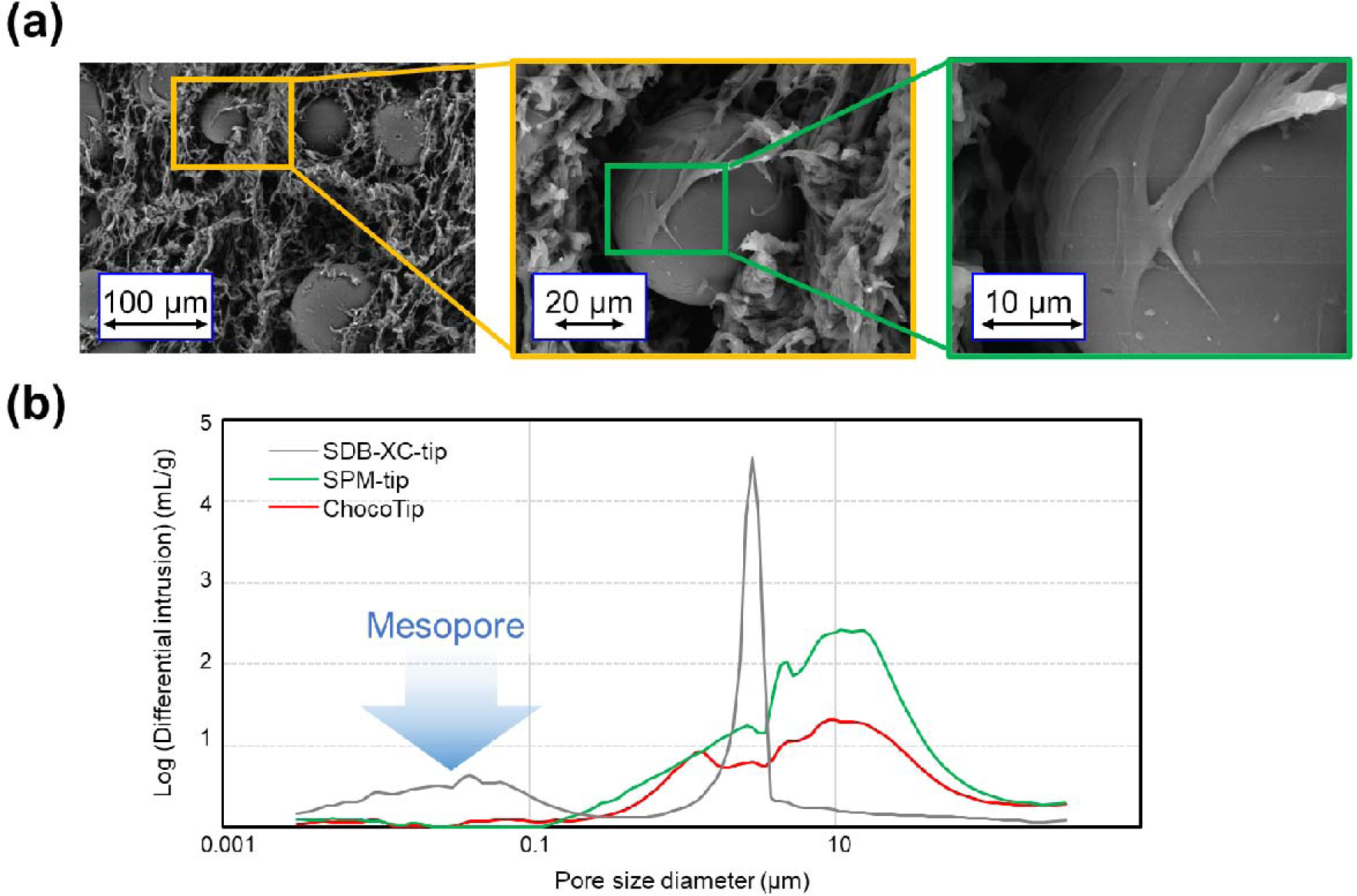
Morphological characterization of ChocoTip. (a) SEM image of the surface of ChocoTip. The surface of the St-DVB particle is partially enveloped by the fibrous structure of SPM. (b) Pore characterization of SDB-XC-tip, SPM-tip, and ChocoTip by mercury porosimetry.

To test this hypothesis, we reduced the number of pieces of Empore SDB-XC disk in the SDB-XC-tip from three to one, anticipating a decrease in irreversible peptide adsorption into the mesopores of St-DVB particles. As expected, the peptide recovery efficiency at shorter retention times was higher with one piece of Empore SDB-XC disk compared to three pieces, suggesting that the lower peptide recovery efficiency in SDB-XC-tip was likely due to irreversible adsorption in the mesopores (Figure S5). However, even with this reduction in the number of pieces of Empore SDB-XC disk, the recovery efficiency of peptides with longer retention times remained higher with ChocoTip. Furthermore, the irreversibly adsorbed peptides in SDB-XC-tip could not be eluted even with strong elution buffers containing higher concentrations of ACN (Figure S6). In another experiment, we prepared a Stacking-SPM-tip by wet-packing 1.0 mg of St-DVB particles into the SPM-tip (Figure S7a). Then, we utilized Stacking-SPM-tip for desalting 20 ng of tryptic peptides to compare the recovery efficiency with that of ChocoTip (Figure S7b). The sum of peptide intensities was much higher in the case of ChocoTip than Stacking-SPM-tip, indicating superior recovery efficiency with ChocoTip. These results strongly support our hypothesis and indicate that the unique morphology of ChocoTip formed in the thermal mixing process is the reason for the significantly reduced sample loss and high recovery in the desalting step using the StageTip method.

## CONCLUSIONS

In this study, we developed ChocoTip as a novel peptide purification technique for MS-based proteomics. The stationary phase of ChocoTip was synthesized by thermally mixing two types of hydrophobic polymers, thermoplastic EVA resin and St-DVB particles. The sponge-like flexibility, derived from the monolithic EVA carrier, simplified the column packing process, offering both ease and reproducibility. Peptide samples were comprehensively retained on the hydrophobic surface of ChocoTip, composed of embedded St-DVB particles and the monolithic carrier, while thermally melted EVA resin effectively prevented irreversible adsorption onto the mesopores of St-DVB particles by filling up these pores. ChocoTip outperformed standard SDB-XC-tip in peptide recovery efficiency, enabling the identification of more than double the number of peptides from 20 ng of tryptic peptides of Hela cell lysates. In addition, the intensities of commonly identified peptides were higher with ChocoTip than with SDB-XC-tip over the entire retention time range, further underscoring the high recovery obtained with ChocoTip. This advantage is particularly pronounced for hydrophobic and longer peptides. Thus, ChocoTip is a promising platform for peptide purification in ultra-sensitive proteomics, paving the way for improved analytical capabilities in a variety of proteomics applications.

## AUTHOR CONTRIBUTIONS

Experiments were designed by E. Kanao, T. Kubo, and Y. Ishihama. LC/MSMS analysis was performed by S. Tanaka, E. Kanao, and A. Tomioka. StageTips were manufactured by S. Tanaka, E. Kanao and T. Tanigawa. The manuscript was written by S. Tanaka, E. Kanao, and Y. Ishihama.

## ACKNOWLEDGEMENTS

We would like to thank members of the Department of Molecular Systems BioAnalysis and Laboratory of Proteomics and Drug Discovery in Kyoto University for fruitful discussions. This work was supported by JST Strategic Basic Research Program CREST (No. JPMJCR1862), AMED-CREST program (No. JP18gm1010010) to YI, NEDO Intensive Support Program for Young Promising Researchers (seeds-4891) to EK, Grants-in-Aid for Scientific Research (KAKENHI; Grants JP23K13774, JP21K14652, and JP20K20567 to EK, 23H04924, 23K18185 and 21H02459 to YI) and Incubation Program Advance of Kyoto University to EK and TK.

## ASSOCIATED CONTENT

### Supporting Information

SEM images of the cross-section surface of SPM-tip and SDB-XC-tip, a comparison of the number of identified peptides binned by retention time between ChocoTip and SDB-XC-tip, distribution of amino acid residues of unique peptides and commonly identified peptides between ChocoTip and SDB-XC-tip, LC/TIMS/Q/TOF analysis of peptides desalted with ChocoTip and SDB-XC-tip, the effect of the number of pieces of Empore SDB-XC disk in StageTip on peptide recovery efficiency, the effect of the ACN concentration in the elution buffer on peptide recovery efficiency with SDB-XC-tip, and desalting performance of Stacking-SPM-tip. This material is available free of charge at http://xxxx.

### Data availability statement

The MS raw data and analysis files have been deposited at the ProteomeXchange Consortium (http://proteomecentral.proteomexchange.org) via the jPOST partner repository (https://jpostdb.org) with the dataset identifier JPST002969.^51^

## Table of contents

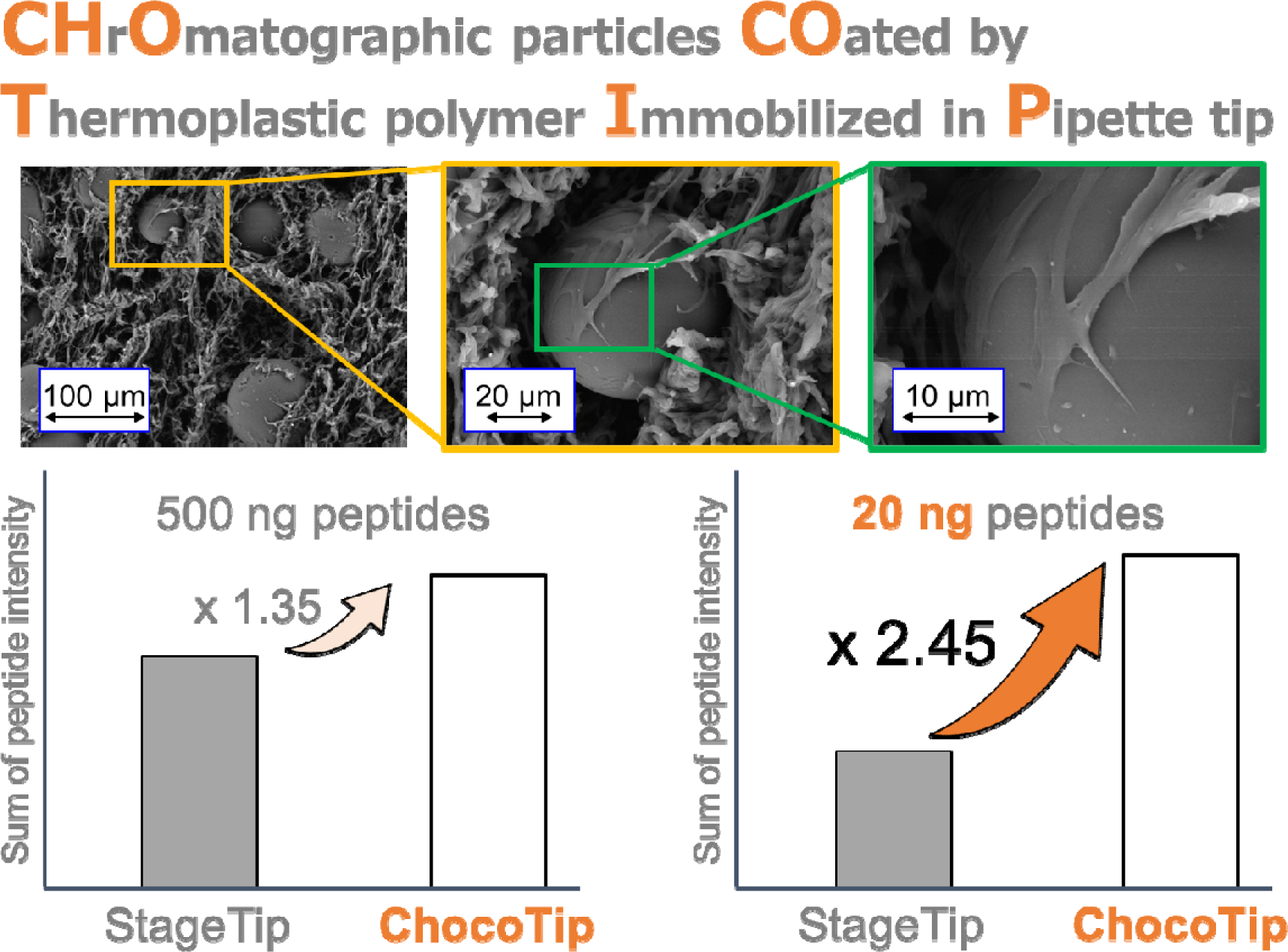

