## Supplemental Information for "High-Recovery Desalting Tip Columns for a Wide Variety of Peptides in Mass Spectrometry-based Proteomics"

**Table of Contents**

Figure S1.	SEM images of the cross-section surface of SPM-tip and SDB-XC-tip.

Figure S2.	Comparison of the number of identified peptides binned by retention time between ChocoTip and SDB-XC-tip.

Figure S3.	Distribution of amino acid residues of unique peptides and commonly identified peptides between ChocoTip and SDB-XC-tip.

Figure S4.	LC/TIMS/Q/TOF analysis of desalted peptides with ChocoTip and SDB-XC-tip.

Figure S5.	Effect of the number of pieces of Empore SDB-XC disk in StageTip on peptide recovery efficiency.

Figure S6.	Effect of ACN concentration in the elution buffer on peptide recovery efficiency with SDB-XC-tip.

Figure S7.	Desalting performance of Stacking-SPM-tip.

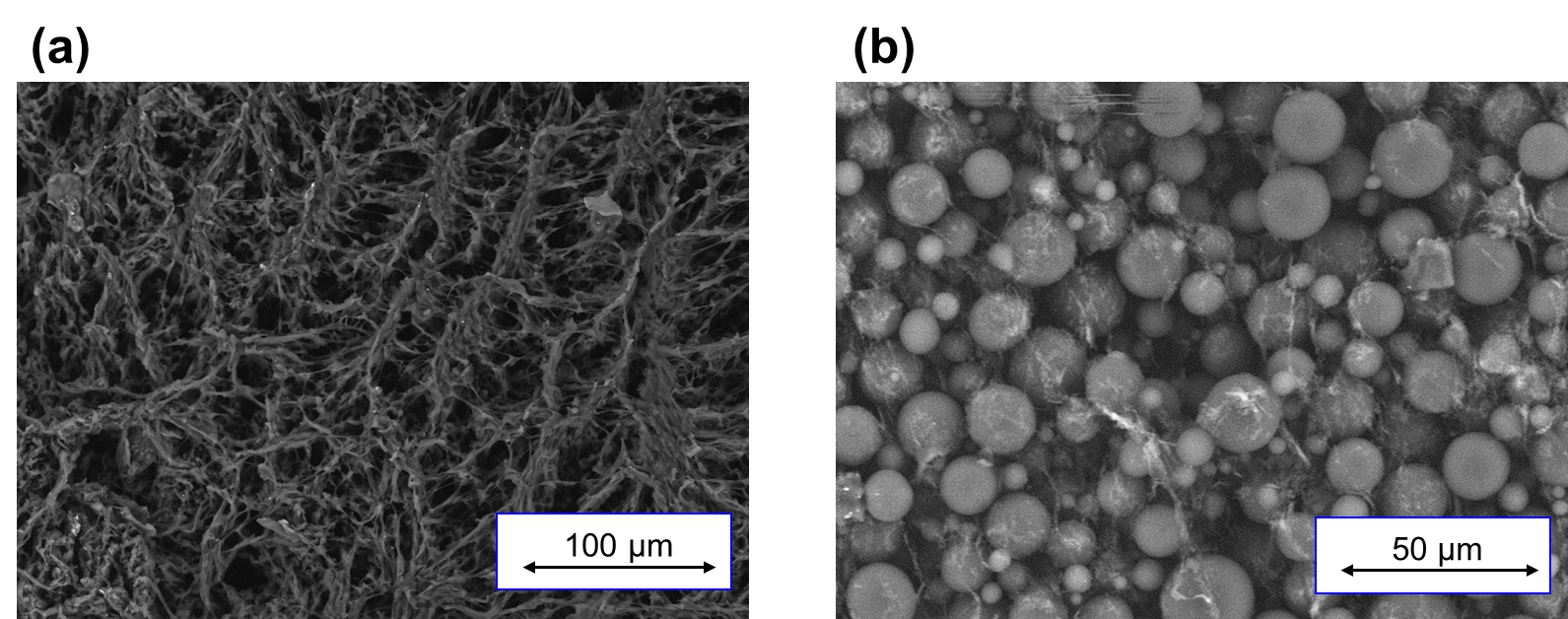

**Figure S1. SEM images of the cross-section surface. (a) SPM-tip, (b) SDB-XC-tip.**

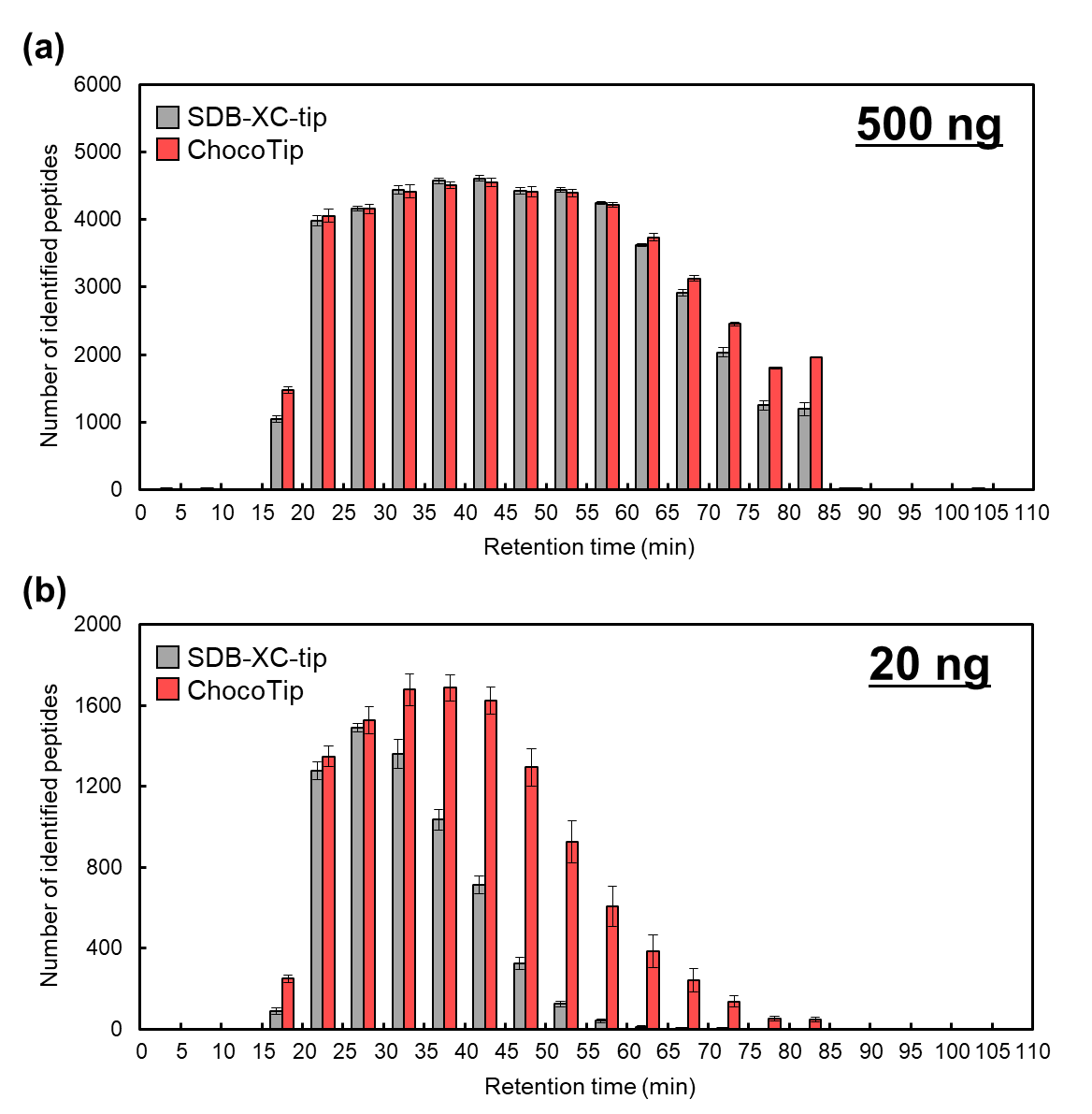

**Figure S2. Comparison of the number of identified peptides binned by retention time between ChocoTip and SDB-XC-tip.** StageTips were used for desalting (a) 500 ng or (b) 20 ng of tryptic peptides from HeLa cell lysates, and a sample equivalent to (a) 250 ng or (b) 10 ng of peptides was injected into the LC/MS/MS system. The error bars indicate the SDs of triplicate analyses with three StageTips.

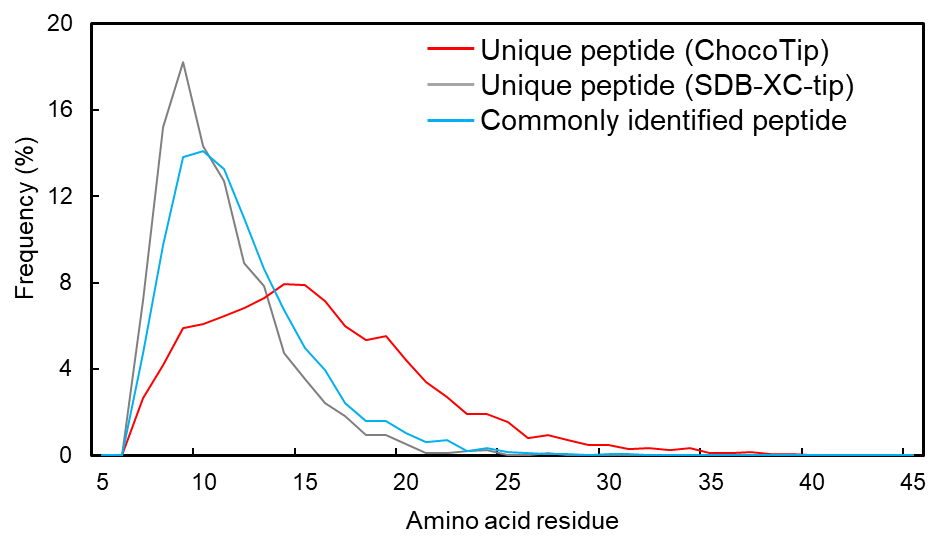

**Figure S3. Distribution of amino acid residues of unique peptides and commonly identified peptides between ChocoTip and SDB-XC-tip.** This data was analyzed using the same datasets as employed for Figure 2b.

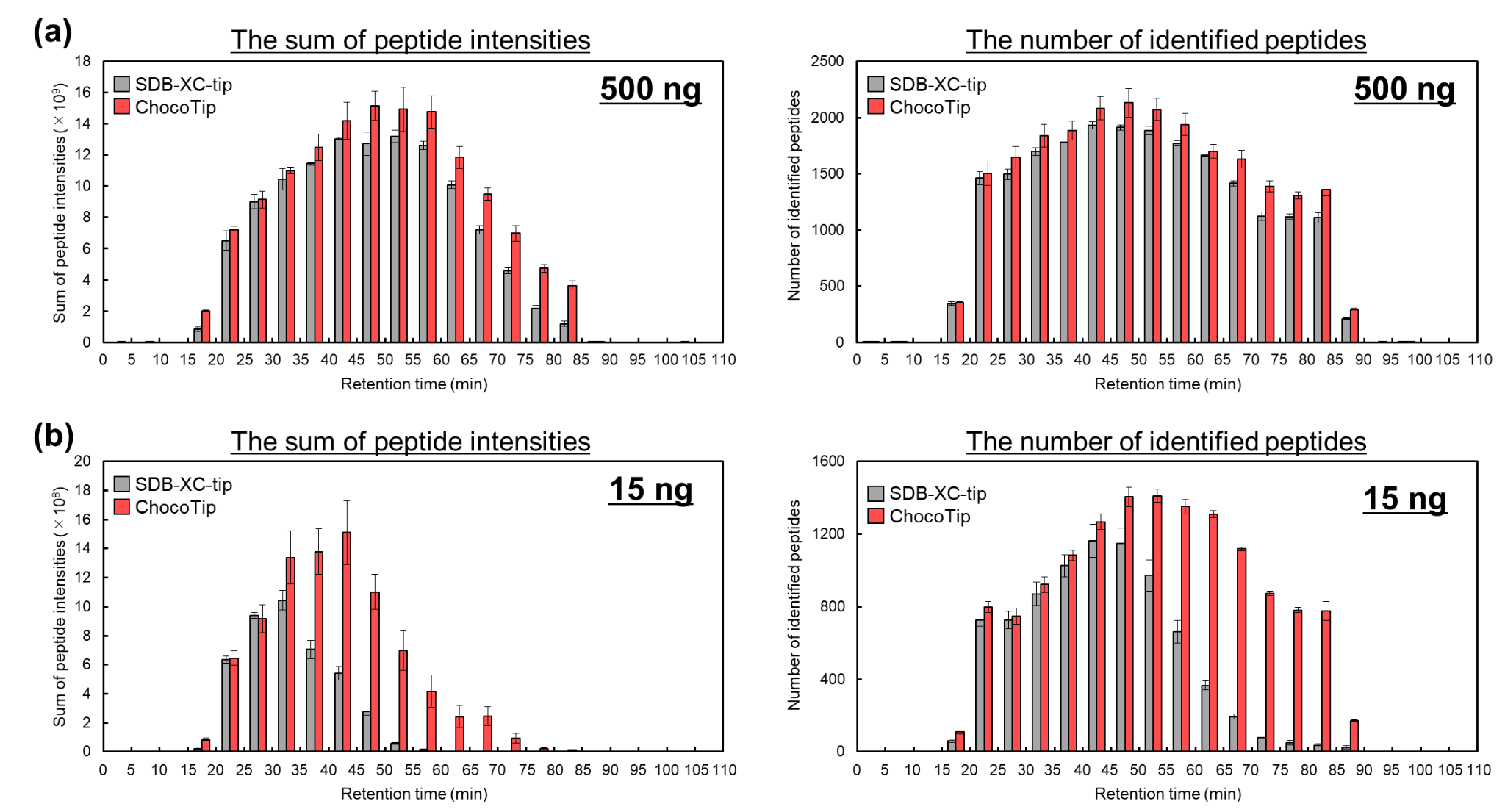

**Figure S4. LC/TIMS/Q/TOF analysis of the desalted peptides with ChocoTip and SDB-XC-tip. The sum of peptide intensities and the number of identified peptides were binned by retention time.** StageTips were used for desalting (a) 500 ng or (b) 15 ng of tryptic peptides from HeLa cell lysates, and a sample equivalent to (a) 250 ng or (b) 10 ng of peptides was injected into the LC/MS/MS system. The error bars indicate the SDs of triplicate analyses with three StageTips.

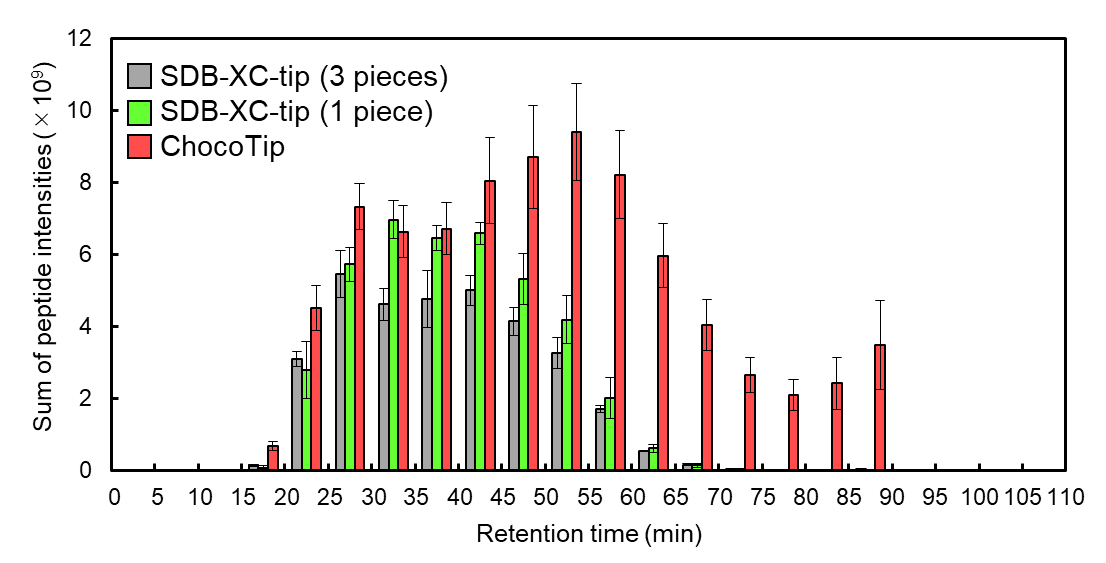

**Figure S5. Effect of the number of pieces of Empore SDB-XC disk in StageTip on peptide recovery efficiency.** The sum of peptide intensity was binned by retention time. StageTips were used for desalting 20 ng of tryptic peptides from HeLa cell lysates, and a sample equivalent to 10 ng of peptides was injected into the LC/MS/MS system. The error bars indicate the SDs of triplicate analyses with three StageTips.

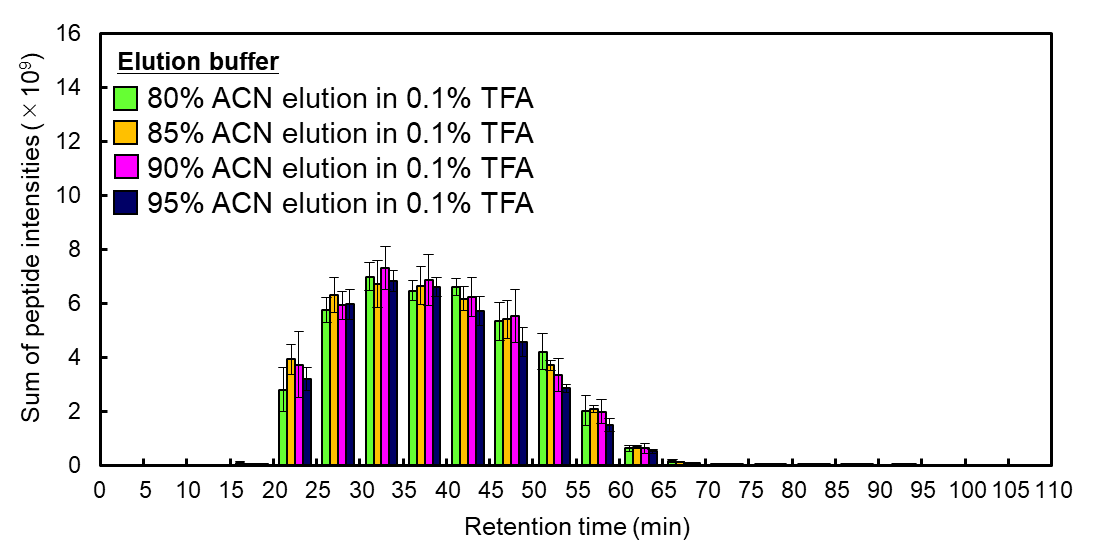

**Figure S6. Effect of ACN concentration in the elution buffer on peptide recovery efficiency with SDB-XC-tip.** SDB-XC-tips were manufactured by stamping out a single piece of the Empore SDB-XC disk with a 16-gauge syringe needle and packing it into a 200 µL pipette tip. The sum of peptide intensity and the number of identified peptides were binned by retention time. SDB-XC-tips were used for desalting 20 ng of tryptic peptides from HeLa cell lysates, and a sample equivalent to 10 ng of peptides was injected into the LC/MS/MS system. The error bars indicate the SDs of triplicate analyses with three SDB-XC-tips.

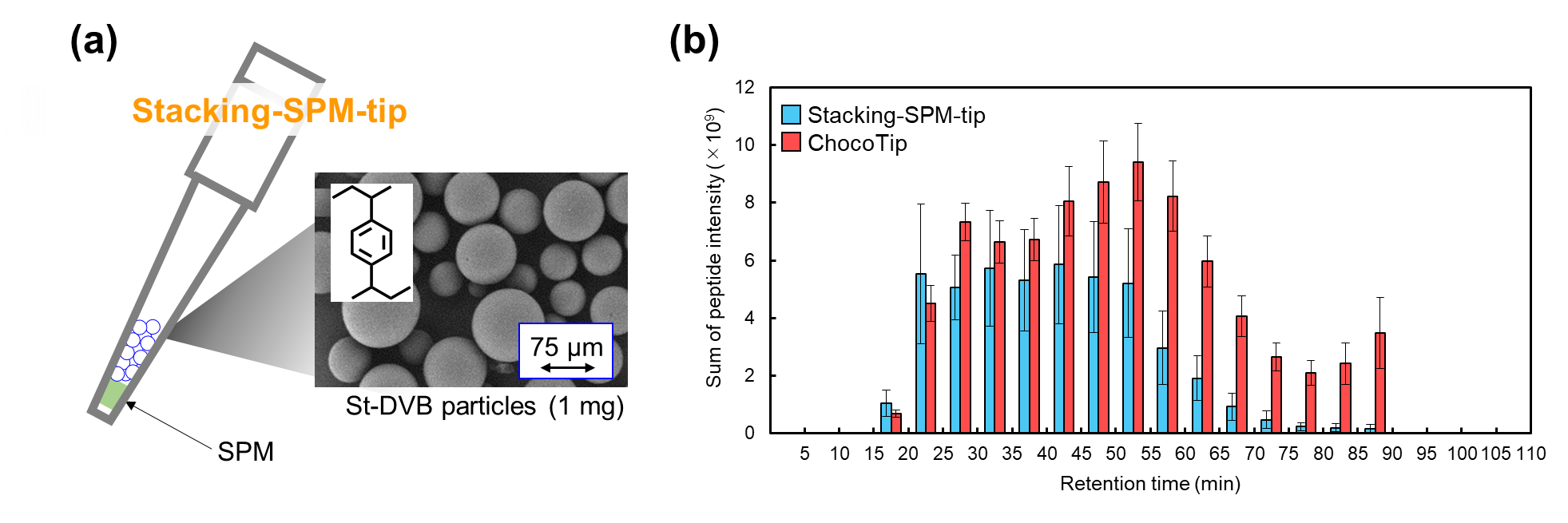

**Figure S7. Desalting performance of Stacking-SPM-tip.** (a) Schematic diagram of stacking-SPM-tip. Stacking-SPM-tip was prepared by wet-packing 1.0 mg of St-DVB particles into SPM-tip. (b) Comparison of the number of identified peptides binned by retention time between ChocoTip and Stacking-SPM-tip. StageTips were used for desalting 20 ng of tryptic peptides from HeLa cell lysates, and a sample equivalent to 10 ng of peptides was injected into the LC/MS/MS system. The error bars indicate the SDs of triplicate analyses with three StageTips.
